# Complete identity of the *cis*-asRNA StfZ and its influence on FtsZ protein level and cell division in *Escherichia coli*

**DOI:** 10.1101/2022.03.08.483438

**Authors:** Deepak Anand, Kishor Jakkala, Rashmi Ravindran Nair, Deepti Sharan, Atul Pradhan, Nagaraja Mukkayyan, Parthasarathi Ajitkumar

## Abstract

Bacteria regulate FtsZ protein levels through transcriptional and translational mechanisms for proper cell division. A *cis*-antisense RNA, StfZ, produced from the *ftsA-ftsZ* intergenic region was proposed to regulate the FtsZ level in *Escherichia coli*. However, its structural identity remained unknown. In the present study, we determined the complete sequence, identified the promoters, and effects of StfZ on the FtsZ level. We show that StfZ is expressed at 1:6 ratio of StfZ:*ftsZ* mRNA at all growth phases from three promoters as three isoforms of 366 nt, 474 nt, and 552 nt. Overexpression of StfZ reduces FtsZ level, increases cell size, and blocks cell division. Thus, the *cis*-encoded StfZ emerges as a novel factor for maintaining the FtsZ level at different growth phases in *E. coli*.

## Introduction

FtsZ is an essential protein for cell division and cytoskeletal integrity in most bacterial cells (Dai and Lutkenhaus, 1991; Pla et al., 1991; Löwe et al., 2004; Shih and Rothfield, 2006). In *E. coli*, the ratio between FtsZ and FtsA molecules (5:1) (Rueda et al., 2003) is important for proper cell division (Ward and Lutkenhaus, 1985; Dai and Lutkenhaus, 1992; Dewar et al., 1992). It was shown that a 2-7 fold increase in FtsZ level results in the mini-cell formation, due to additional division events, whereas further higher level or lower than normal level cause filamentation due to imbalance in cell division machinery (Bi and Lutkenhaus, 1990; Wang and Gayda, 1990). Thus, regulation of the FtsZ level is crucial for proper cell division. The regulation happens at the transcriptional and translational level (Aldea et al., 1990; Cam et al., 1996; Flärdh et al., 1997; Flynn et al., 2003; Tamura et al., 2006; Camberg et al., 2009).

Trans-acting small non-coding asRNAs are usually encoded in intergenic regions on the chromosome and control translation or degradation of their target mRNAs. Generally, each trans-acting non-coding asRNA has multiple target mRNAs and binds near the ribosomal binding site of the target mRNAs (Saberi et al., 2016). One such asRNA, DicF, against *ftsZ* has been found to influence FtsZ protein level in certain strains of *Escherichia coli* (Tétart and Bouché, 1992). Besides DicF, Dewar and Donachie had proposed expression of StfZ *cis*-asRNA from the 60-bp spacer sequence into the beginning of *ftsZ*, blocks cell division when placed in a high copy number plasmid at 42°C. A structural change in the non-coding RNA occurs through binding to small metabolites (riboswitches) or through a change of temperature (thermoregulators) or pH (pH sensors) (Saberi et al., 2016). In both cases, elevated temperature caused the phenotypic effects of asRNA. Also, StfZ was not studied further for its expression and promoter analysis. Since regulation of principal cell division gene FtsZ is an important aspect of cell division, it is important to elucidate its mode of action, and physiological role in cell division.

The present study establishes the complete sequence of StfZ, its growth dependent expression, the stoichiometry of expression against *ftsZ* mRNA, and its role in cell division regulation. We showed that StfZ RNA has three isoforms and identified the respective promoters. Further, we investigated its influence on FtsZ level and thereby on cell division. The generation of a knockout or deletion mutant of the *stfZ* was not possible as the region of interest is shared by *ftsZ* sequence. Nevertheless, the observations reported in the present study show the expression of natural antisense RNA isoforms of StfZ as a novel factor that *E. coli* uses to influence FtsZ level and thereby cell division.

## Materials and Methods

### Bacterial strains, plasmids, and growth

Bacterial strains and plasmids are listed in S1 Table and S2 Table respectively. All the strains were cultured in Luria-Bertani broth or agar for growth. Strains with plasmids were selected on ampicillin (100 µg/ml) or kanamycin (25 µg/ml). For StfZ induction cultures were grown at 25° and shifted to 30°C, 37°C or 42°C as per the experiment.

### cDNA preparation

RNA was isolated using hot-phenol method (Wecker, 1959; Roy et al., 2004). In brief, cells were lysed in lysis buffer (S3 Table). The aqueous phase of RNA was extracted with hot phenol (65°C, pH 5.2) followed by phenol: chloroform and chloroform extractions. RNA was precipitated and dissolved in RNase-free water. RNA preparations were treated with DNase-I which was verified using PCR for a 16S rRNA gene (Condon et al., 1995). cDNA was prepared using 5 µg total RNA with RevertAid-Premium Reverse Transcriptase kit (Fermentas). For each reaction, 20 pmoles of gene-specific reverse primer were added and annealed at 55°C for 10 min, followed by addition of reverse transcriptase for an extension at 55°C for 60 min. The reaction was stopped by incubating at 85°C for 10 min. The cDNA preparation was used for RT-PCR and quantitative PCR.

### RT-PCR and quantitative real-time PCR

RevertAid-Premium Reverse-Transcriptase kit and Evagreen real-time PCR master mix (GBiosciences) were used for RT-PCR and real-time PCR, respectively (Wang et al., 2006). Primers ODA-01 and ODA-02 for StfZ, ODA-03 and ODA-04 for 16S rRNA, ODA-05 and ODA-06 for *ftsZ*, ODA-09 and ODA-10 for *mutgfp* and ODA-11 with ODA-12 for *ftsA* (S4 Table) were used. Reactions were performed as per described protocols. The cDNA of 16S rRNA was used as the normalisation control (Condon et al., 1995). Real-time PCR was performed in Applied Biosystems-ViiA7. The 2^−ΔΔCt^ method was used for quantitation (Livak and Schmittgen, 2001; Giangrossi et al., 2010). The levels of expression were presented as fold changes to the control sample.

### Primer extension assay (PEA)

PEA was performed using 30 µg of total RNA isolated from *E. coli* K12 cells of 0.3 OD_600 nm_ (OD). Primers, ODA-07 and ODA-08, were radiolabeled with [γ-^32^P]-ATP using T4-polynucleotide kinase kit (Fermentas). Labelled primers were purified with Sephadex G-50 column, annealed to RNA and extended at 55°C with 200 U of RevertAid-Premium Reverse-transcriptase for 60 min. Primer extension products were denatured at 95°C and fractionated on 8% polyacrylamide gel containing 7 M urea. A parallel manual sequencing reaction was performed using CycleReader™ DNA Sequencing Kit (Fermentas) and loaded in the lane next to PE reaction. The PCR product template for sequencing was generated using primer ODA-13 in combination with ODA-08 or ODA-07 (S4 Table). Autoradiography was performed using phosphorimager after 24 hrs of exposure of the sample.

### Molecular cloning

Region of *stfZ* was amplified form *E. coli* genomic DNA using primer ODA-14 and ODA-15. PCR product was digested with KpnI and XabI and ligate to pBSKS vector at same sites.

The StfZ promoters were cloned using oligonucleotide annealing method (Arumugam et al., 2012). Oligonucleotides for P1 (ODA-16 and ODA-17), P2 (ODA-18 and ODA-19), P3 (ODA-20 and ODA-21) and the respective -10 deletion P1 (ODA-22 and ODA-23), P2 (ODA-24 and ODA-25), and P3 (ODA-26 and ODA-27) were annealed and cloned at KpnI and BamHI site in pFPV27 (Valdivia and Falkow, 1996). The promoter regions from P1 to P3 were amplified using ODA-28 and ODA-29 (S4 Table) and cloned at KpnI & BamHI sites in pFPV27. A region from 1098^th^ nt of *ftsA* to 197^th^ nt of *ftsZ* gene was amplified using ODA-14 and ODA-15 primers (S4 Table) and cloned at the KpnI/XbaI sites in pBS(KS) (Alting-Mees and Short, 1989) to get PAK12 strain.

### 3′ RACE

Twenty micrograms of total RNA from 0.5 OD_600 nm_ culture was enriched for total mRNA using Ribominus™ Transcriptome Isolation kit (Invitrogen K1550-03). The 5′ phosphorylated ODA-30 oligo (S4 Table) was ligated to the 3′ ends of ribominus RNA using T4 RNA ligase. The cDNA was synthesised with the complementary oligo ODA-31. StfZ-specific cDNA was PCR amplified using ODA-01 and ODA-31 primers, using standard conditions but at the appropriate temperature of annealing of the primers. The PCR product, extracted from agarose gel, ethanol precipitated, washed, and was cloned in pBS(KS) and sequenced.

### Northern hybridization

Biotin Labelled DNA probe was synthesised using PCR amplified *stfZ* region from plasmid pDA09. First, primer ODA-14 and ODA-015 were used with Taq DNA polymerase (Thermoscientific) then 100 ng of PCR product with 0.5 µM of ODA-15 primer, 0.05 mM each of dATP, dCTP, dGTP and biotin-11-dUTP (Thermoscientific) was used. Twenty such reactions were pooled and precipitated with 3 M ammonium acetate and ethanol. DNA probes were dissolved in DNase free water. RNA probe was generated against *stfZ* region by *in vitro* transcription from KpnI digested pDA9 plasmid. HiScribe™ T7 High Yield RNA synthesis kit (NEB) was used as per the manufacturer’s protocol with Biotin Labelling RNA Mix (Roche) to obtain *stfZ* complementary probe.

For northern hybridization, RNA from PAK02 and PAK12 strains (0.2 ODD) were fractionated on 10% polyacrylamide denaturing gel with 7 M urea. RNA was blotted to a positively charged nylon membrane (Roche). The membrane was subjected to UV cross-linking (1200 µJ/cm^2^ for 20 min) and dried at 50°C for 30 min. The membrane was blocked in a pre-hybridisation buffer (Table 3) at 60°C for 3 hrs. Pre-hybridisation buffer was replaced with hybridisation buffer containing biotin-labelled DNA probe (denatured at 95°C for 5 min and snap-chilled on ice) and incubated overnight at 60°C. The nylon membrane was washed thrice with wash buffer (1x SSC containing 0.1% SDS) for 15 min each, at room temperature. The membrane was then blocked for 15 min and incubated with streptavidin-HRP conjugate (Genei) (1:2000 dilution) for 15 min. The membrane was washed and developed using Clarity Western ECL Substrate (Bio-Rad). The same protocol was followed in the case of the control sample except that 160 µg of total RNA, isolated from 0.5 OD_600 nm_ PAK02 culture, was used for northern blotting and the probe used was 10 µg of biotin-labelled RNA obtained from *in vitro* transcription.

### Immunofluorescence microscopy

Immunostaining was performed as described (Addinall et al., 1996), with a few modifications. The harvested cells were fixed with 0.4% paraformaldehyde and 0.25% glutaraldehyde solution for 10 min at room temperature and 50 min on ice. The cells were washed with 1x PBS (pH 7.4) (S3 Table), layered over poly-L-lysine (0.1%, w/v) coated multi-well slide, permeabilised with 2 mg/ml of lysozyme (Sigma), blocked with BSA (2% w/v in PBS), followed by incubation with 1:500 dilution of affinity-purified rabbit polyclonal anti-FtsZ antibody overnight at 4°C in a humid chamber. The cells were washed five times with 1x PBS, followed by 60 min incubation with 1:1000 dilution of Cy3 anti-rabbit IgG antibody (0.1 µg/ml; Sigma). The cells were washed again with 1x PBST (S3 Table) and incubated with 0.5 µg/ml DAPI for 5 min. DAPI was washed off with 1x PBST solution and cells were mounted with 80% glycerol. Images were taken under the Zeiss AxioImager M1 fluorescence microscope. AxioVision software was used for size measurements and image processing.

### Western blotting

Total protein (100 µg) was fractionated on 10% polyacrylamide gel and blotted onto methanol activated PVDF membrane. PVDF membrane was blocked overnight in blocking buffer at 4°C. The blocking buffer was replaced with rabbit anti-FtsZ primary antibody (1:10000) solution for FtsZ. The membrane blots were washed and incubated with 1:10000 diluted anti-rabbit goat IgG (Sigma) (Srivastava et al., 2016). The blots were washed and developed with x-ray film (Kodak) or chemiluminescence imager (ImageQuant LAS 4000) using enhanced luminol reagent (Sigma).

### SYTO9/ PI staining

SYTO9 and propidium iodide (PI) were prepared according to manufacturer’s instructions (Live/Dead Bacterial Kit, Molecular Probes, (Brantl, 2007)). Cells in 50 µl of culture were stained with 0.1 mM SYTO9/PI for 15 min in the dark at 25°C, washed with PBS solution, and images were taken with excitation and emission of 483/503 nm for SYTO9 and 485/630 nm for PI.

## Results

### StfZ RNA is expressed at all growth phases

We determined the presence of StfZ transcript in *E. coli* K12 cells from 0.2 (early log phase) to 2.5 (late stationary phase) OD_600 nm_ (hereinafter called OD) cultures. RT-PCR analysis, performed using ODA1 and ODA-02 primers located across the intergenic region of *ftsA* and *ftsZ*, showed the presence of 177 bp RT-PCR product from all the growth phases (Fig. 1A). Cloning and sequencing of the StfZ RT-PCR product confirmed its synthesis from the complementary strand in the *ftsA*-*ftsZ* intergenic region. These observations suggested that the StfZ is transcribed from the *ftsA*-*ftsZ* intergenic region at all growth phases.

**Figure. 1.**
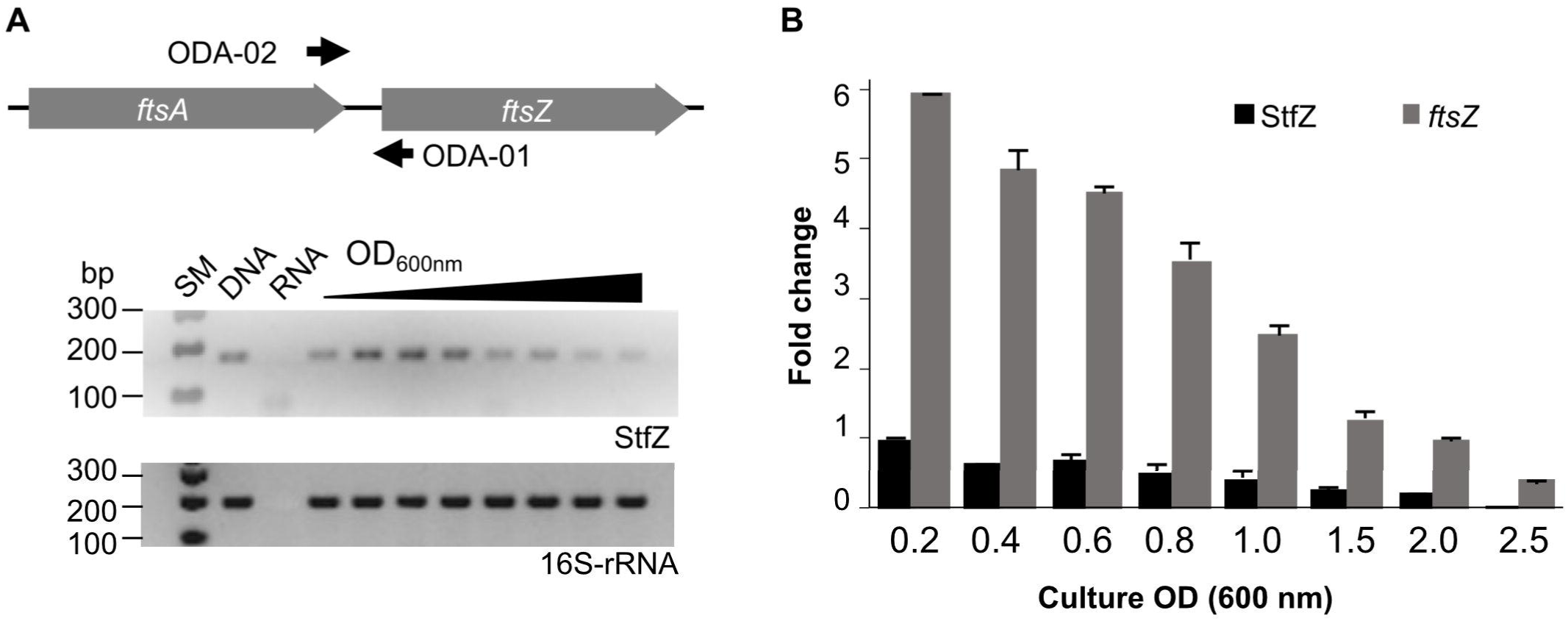
Detection and quantitation of StfZ RNA. RT-PCR for the detection of StfZ transcript. (**A**) Upper panel shows the location of the primers ODA-01 and ODA-02 used for StfZ RT-PCR. Oligo ODA-02 was used for StfZ cDNA synthesis. The lower panel shows the RT-PCR products of StfZ and 16S rRNA. G-DNA, genomic DNA as the positive control; RNA, RNA sample from 0.6 OD as the negative control; and cDNA from 0.2 to 2.5 OD cultures. The RT-PCR products were of 177 bp (StfZ) and 218 bp (16S-rRNA). (**B**) Semi-quantitative real-time PCR of *ftsZ* mRNA and StfZ transcript from *E. coli* K12 cells from 0.2 to 2.5 OD cultures. *ftsZ* cDNA was synthesised with ODA-06 oligo and StfZ cDNA using ODA-02 oligo. The bar graph represents the levels of the transcripts with respect to StfZ at 0.2 OD.

### Stoichiometric expression of StfZ RNA to *ftsZ* mRNA

To find the stoichiometry between antisense StfZ RNA and its target sense RNA, the *ftsZ* mRNA, at different growth phases, both RNA were analysed using semi-quantitative RT-PCR. The StfZ RNA level was highest at the early log phase (0.2 OD) and was maintained at a relatively high level till the mid-log phase (0.6 OD) (Fig. 1B). The StfZ RNA levels progressively decreased as the culture approached the stationary phase (1.5 OD), where the level was about 5-fold lower compared to 0.2 OD samples. Subsequently, the level further reduced at 2.5 OD (Fig. 1B). The decrease in the StfZ RNA level was found to be growth dependent. The reduction in the level of StfZ RNA correlated with the steady decrease in the level of its target, *ftsZ* mRNA (Fig. 1B). Consistent with this correlation, the ratio between StfZ RNA to *ftsZ* mRNA was found to be always ∼1:6, irrespective of the growth phase. This indicated a coordinated expression of the sense and the antisense RNAs in a growth phase-dependent manner.

### Multiple transcripts of StfZ

For determining the 5′ and 3′ ends and size of StfZ RNA, primer extension analysis (PEA), 3′ rapid amplification of cDNA ends (3′ RACE) and northern hybridisation were performed. PEA was performed using ODA-07 or ODA-08 (S4 Table, Fig. 2A) primers located in the intergenic region of *ftsA* and *ftsZ*. We obtained two products from the extension of the ODA-07 primer. The first product was located at 117^th^ and the second at 195^th^ positions downstream to *ftsZ* ATG start codon (Fig. 2B). Primer ODA-08 positioned downstream to the ODA-07 binding site, produced three products (Fig. 2C). The first two products were overlapping with the product obtained from ODA-07 while the additional product was at the 9^th^ position downstream to the *ftsZ* start codon. These three PE products were named TSS-9, TSS-117, and TSS-195, according to the distance from *ftsZ* 5′ end. Subsequently, we compared the consensus sequences of -10 and -35 region of *E. coli* promoters (Hawley and Mcclure, 1983; Lisser and Margalit, 1993; Mitchell et al., 2003) with the sequence in the region upstream of the 5′ end nt of the three respective PE products. Thus, the predicted promoter sequences for TSS-9, TSS-117, and TSS-195 were named P1, P2 and P3, respectively (Fig. 2D). P1 and P2 showed -10 consensus with TATAAT and -35 consensus with TTGACA of the experimentally identified promoters of *E. coli* (Hawley and Mcclure, 1983; Lisser and Margalit, 1993; Hershberg et al., 2001; Mitchell et al., 2003). The predicted -10 and -35 sequences for the putative P3 promoter showed divergence (Fig. 2D). Nevertheless, the -35 sequence (CCAACTT) of P3 promoter showed consensus with the -35 sequence (GGAACTT) of *rpoEp*3 gene of *Salmonella enterica serovar Typhimurium* (Skovierova et al., 2006). It has been demonstrated that many of the promoters of antisense RNAs of enteric bacteria do not show conservation in the -10 and -35 sequences (Raghavan et al., 2012). The putative P3 promoter might be one such promoter.

**Figure. 2.**
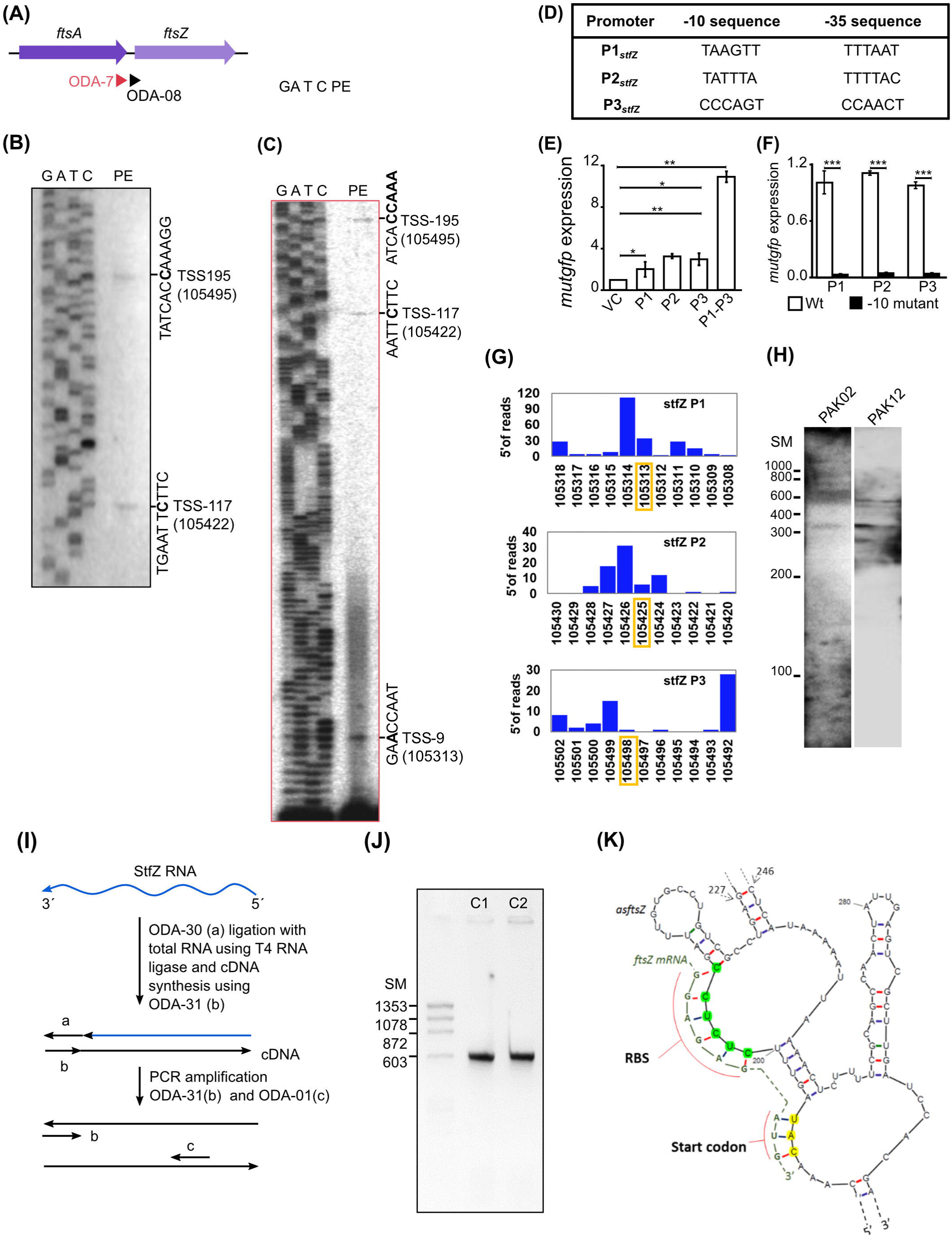
Identification of 5′ end, promoter and 3′ end of StfZ. (**A**) Schematic representation of the Primer extension analysis (PEA) products showing the positions of the oligos used for the extension. (**B, C**) PEA radiograph for StfZ RNA. (**B**) PEA with oligo ODA-07. Lane GATAC are the sequence product lanes corresponds to ddG, ddA, ddT and ddC termination reactions. Lane PE, primer extension product. **(C)** PEA with oligo ODA-08. (**D**) Putative promoters of the *stfZ* gene with their respective -10 and -35 sequences predicted with respect to the 5′ ends of the three PEA products. (**E**) Promoter assay using *mutgfp* as a reporter in pFPV27 vector. P1, P2 and P3 predicted promoters with native sequence (wt) (white bars) and respective -10 deletion mutant -10 mutant (black bars) were cloned upstream to *mutgfp*. Strain PAK05 (P1 wt), PAK07 (P2 wt), PAK09 (P3 wt), PAK06 (P1Δ-10), PAK08 (P2Δ-10) and PAK10 (P3Δ-10). Y-axis indicates the relative expresion (**F**) Relative activity of predicted *stfZ* promoters, P1, P2, and P3, individually and cumulative were analysed in PAK05, PAK07, PAK09 and PAK11 strains respectively. pFPV27, containing *mutgfp* without promoter was used as the vector control. Statistical analysis is indicated with asterisks (*p* < 0.05 *, *p* < 0.01 **, *p* < 0.001 ***). (**G**) Transcription start site (TSS) frequency bar-plot from existing RNA seq data represents the surrounding region of StfZ 5′ end identified by primer extension. *stfZ*-P1, *stfZ*-P2 and *stfZ*-P3 in this graph represent the promoters with respect to TSS-9, TSS-117 and TSS-195. The y-axis on each graph represents the number of reads starting on each position and x-axis represents the genomic positions. RNA-Seq data was extracted from NCBI-SRA (accession number-SRX3413960). The orange box shows the TSS determined from PEA in this study. (**H**) Northern blot for PAK02 and PAK12 RNA samples. PAK02 and PAK12 were probed with single-stranded RNA probe. RNA ladder was used as a size marker. The asterisks indicate three bands of expected sizes, 366 nt, 474 nt, and 552 nt, approximately. (**I**) Strategy for 3′ RACE to identify the 3′ end of StfZ RNA. (**J**) Approximate size of the 3′ RACE product is 600 bp. (**K**) 5′ Region of the predicted secondary structure of StfZ RNA from Mfold web server. The *ftsZ* (yellow) ribosomal binding site (RBS) and of the AUG start codon complementary to the StfZ structure are shown next to the open loops, A and B, respectively.

### The predicted promoters of StfZ RNA drive reporter gene expression

The -10 regions of bacterial promoters are crucial for the initial stages of sigma factor interaction. Transcription initiation drastically fails in the absence of -10 element (Ruff et al., 2015; Browning and Busby, 2016). This characteristic feature has been used to validate and map bacterial promoters. Taking the same approach to validate the predicted promoters, we constructed Δ-10 promoter constructs (pDA3, pDA5, and pDA7) (S2 Table), with *mutgfp* as the reporter gene, and compared its expression from respective native promoter constructs (pDA2, pDA4, and pDA6) (S2 Table). These constructs were expressed from JM109 strain (S1 Table). Promoter activity was quantitated using real-time PCR for *mutgfp* mRNA with ODA-09 and ODA-10 primers (S4 Table). cDNAs for *mutgfp* were synthesised using primer ODA-10 from total RNA isolated from the promoter construct transformants grown to the mid-log phase (0.6 OD).

Deletion of predicted -10 elements of all the putative promoters showed about 20-fold reduced *mutgfp* expression (Fig. 2E). The transcriptional activity of the three predicted promoters and its abrogation in the -10 deletion mutants validated the authenticity of the promoters. Relative expression of the putative promoters showed a different level of expression, while a significant level of cumulative expression was observed from the combined P1-P2-P3 promoter construct (Fig. 2F). These observations, in turn, implied transcription of *stfZ* from three independent promoters producing three isoforms of StfZ. Interestingly, the P1 promoter region corresponded to the previously predicted promoter for StfZ (Dewar and Donachie, 1993).

Northern blot against StfZ was performed from *E. coli* K12 (wt) and PAK12 transformant carrying cloned portion of *stfZ* gene (S1 Table), showed bands in the range of ∼350, ∼450 and ∼500 nts (Fig. 2H). These bands corresponded to the sizes of the PEA products (see Fig. 2C). The presence of additional bands in the northern blot could be the result of non-specific hybridisation. The consistent presence of the expected three bands despite high stringency washes with 0.1x SCC and 0.1% SDS at 55°C indicated their authenticity. The very low intensity of the bands suggested a low level of StfZ expression, which could be observed by RT-PCR only (see Fig. 1A).

In addition to our experimental analysis, we examined the existing RNA-seq data to know whether StfZ transcripts were detected before this study. RNA-Seq data were extracted from NCBI SRA database (accession number - SRX3413960) (Livingstone et al., 2018) and analysed for TSS frequency per nucleotide in *stfZ* region. TSS frequency was plotted as bar graphs along the DNA sequence. The TSS of StfZ were detected, wherefrom TSS-9, TSS-117 and TSS-195 RNAs are transcribed (Fig. 2G). However, the detection does not match precisely in the RNA-seq +1 TSS frequency.

Additionally, to verify the exact ends of the isomers and to find out if it is a processed RNA product, we performed circular RACE with and without Tobacco acid pyrophosphatase (TAP) treatment of RNA. TAP removes 5′ cap of RNA therefore a processed RNA can be detected in circular RACE without TAP treatment but not a capped RNA. Following this method (McGrath, 2011) we did not find any amplification from StfZ region and any condition.

### StfZ RNA 3′ end extends to upstream of *ftsZ*

After determining the 5′ ends of StfZ transcripts, we performed 3′ RACE to find out the respective 3′ ends and thereby the exact size of the transcripts. For this, the ribominus RNA fraction (devoid of ribosomal RNAs) was ligated to the adaptor oligo, ODA-30, and the cDNAs were synthesised using ODA-31, as indicated in the cartoon (Fig. 2I; S4 Table). The cDNA product was amplified with ODA-31 and ODA-01 primers to get the PCR product of ∼600 bp (Figs. 2I, J). Replicates of PCR amplified products (∼600 bp) were cloned in plasmid pDA1 and sequenced (Fig. 2J; S2 Table). Sequencing showed 357^th^ nt upstream of *ftsZ* as the 3′ end of all the isoforms. The entire sequence of the three isoforms encompassed the SD sequence of *ftsZ*, entire *ftsA-ftsZ* intergenic region and 297 nts on the 3′ end of *ftsA* gene. From the three different transcription initiation sites, StfZ isoforms are produced as 366 nt, 474 nt, and 552 nt long RNAs. The sequences were deposited in Bankit database with accession numbers; stfZ_366 KX852304, stfZ_474 KX852303 and stfZ_552 KX852302.

### Features of StfZ RNA

Predicted Mfold secondary structure StfZ showed the presence of two successive loops A and B with 5′ CUCUCC 3′, (complementary region of *ftsZ* mRNA RBS, 5′ GGAGAG 3′) and 5′ CAU 3′ (complementary to *ftsZ* initiation codon 5′ AUG 3′) (Fig. 2K). This could facilitate its initial interaction with target mRNA at the RBS site to form a ‘kissing complex’ followed by extension in both directions to form a complete stable duplex (Gerhart et al., 1994; Lease and Woodson, 2004; Brantl, 2007). The StfZ sequence also have 5′ AATAATA 3′ sequence, which resembled the potential consensus sequence for the binding of Hfq (5′ AAYAAYAA 3′) (Lorenz et al., 2010). It is located 156 nt from 3′ end StfZ transcript and it shares complementary region of *ftsA*. The presence of multiple stop codons, 6 in the reading frame 1, 11 each in the reading frames 2 and 3, in the primary sequence of StfZ ruled out the possibility of coding for any short peptide, unlike the possibility predicted in the earlier study (Dewar and Donachie, 1993).

### StfZ RNA targets *ftsZ* mRNA

StfZ sequence features revealed that it can function as an antisense RNA specific to *ftsZ* mRNA. Therefore, the effect of StfZ RNA overexpression on *ftsZ*-*yfp* translation was tested to verify its target specificity and to document the physiological changes brought about by StfZ overexpression. This method allowed measurement of the effect of the antisense RNA against its target, *ftsZ* mRNA by fluorescence microscopy or directly by YFP fluorescence from bacterial cells. In principle, binding of StfZ RNA to *ftsZ-yfp* should rescue FtsZ-YFP overexpression phenotype (cell-filamentation) and reducing YFP fluorescence (Fig. 3A). The culture was induced at 0.6 OD with either 0.1% arabinose (for *ftsZ-yfp* mRNA) or 1 mM IPTG (for *StfZ* RNA) or with both the inducers simultaneously. Expression of FtsZ-YFP was measured at fluorescence level in the culture and at single cell level my microscopy. Multiple FtsZ-YFP rings and a high level of YFP fluorescence were observed in the arabinose-induced cells due to overexpression of FtsZ-YFP (Fig. 3B and 3C.1). A high level of FtsZ-YFP interfered with the division process and induced cell filamentation (Fig. 3C.2). The co-induction of StfZ RNA along with *ftsZ* mRNA significantly rescued the cells from elongation (Fig. 3C.2). Inhibition of *ftsZ-yfp* translation could be inferred from the reduction in the YFP fluorescence level in the FtsZ-YFP induced cells (Fig. 3C.1). These results suggested that StfZ RNA is specific to the sense *ftsZ* mRNA. There is a functional overlap between DicF and StfZ RNAs as they share the target region near the *ftsZ* RBS sequence (Fig. 3D). However, *E. coli* K12 and JM109 strains did not contain DicF RNA as found using RT-PCR (data not shown), ruling out any interference by DicF RNA in the experiments.

**Figure. 3.**
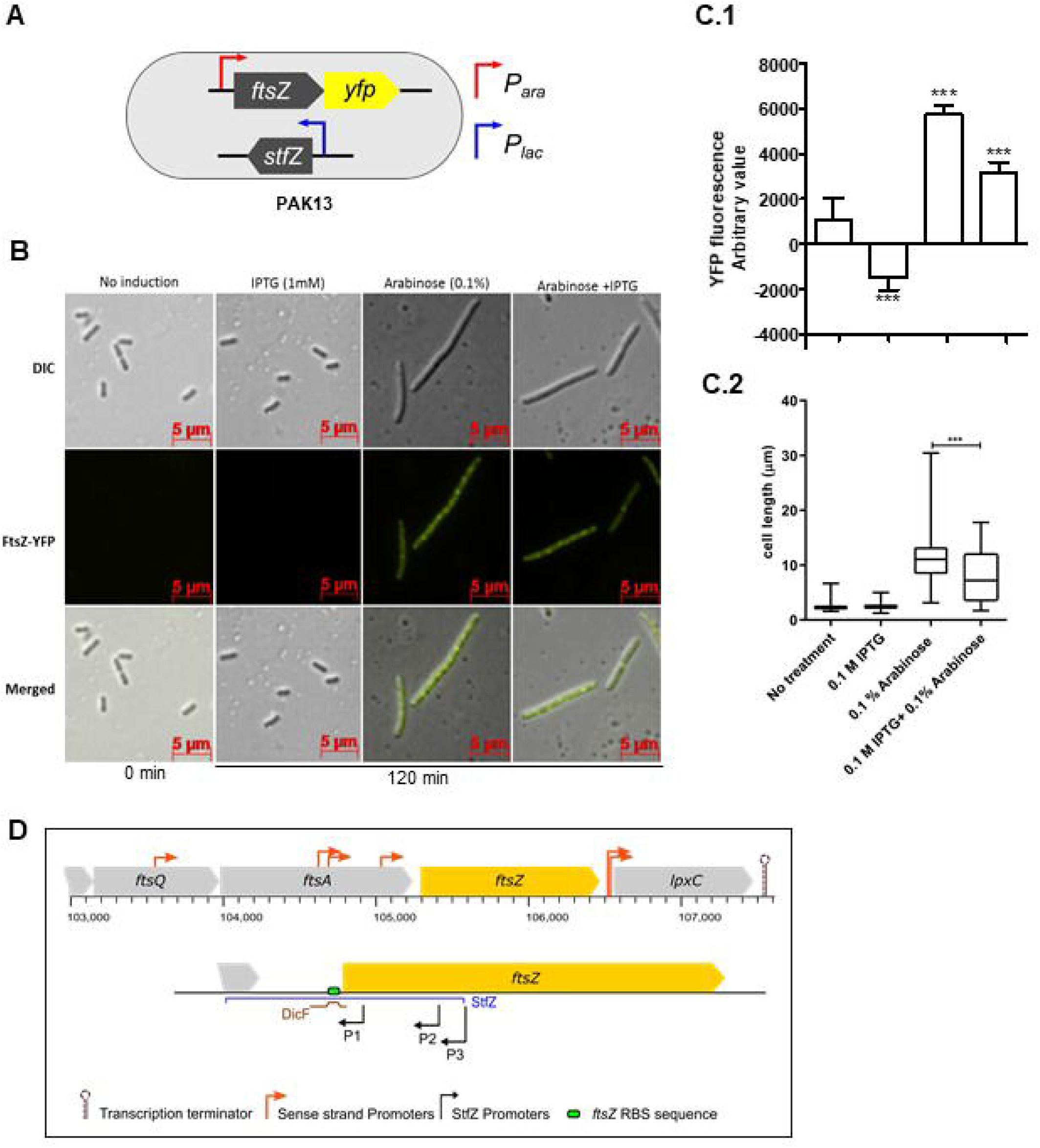
Inhibition of *ftsZ-yfp* mRNA translation by StfZ. PAK13 cells were induced with either 1 mM IPTG or/and 0.1% arabinose at 37°C for 120 min. (**A**) Schematic presentation of PAK13 strain with *P*_*ara*_*-ftsZ-yfp* and *P*_*lac*_*-stfZ* constructs. (**B**) Cells were imaged before induction and after induction for 120 min. Columns: (**a**) negative control without induction of the culture; (**b**) positive control for StfZ RNA expression, induced with IPTG; (**c**) positive control for FtsZ-YFP expression, induced with arabinose; (**d**) experimental sample, induced for the co-expression of StfZ RNA and *ftsZ*-*yfp* fusion mRNA. Top row: DIC images; middle row: FtsZ-YFP images and lower row: merged images. (**C.1**) Quantitation of the YFP fluorescence as arbitrary values for 0 min and120 min post-induction from 200 µl culture. **(C.2)** Box chart for cell-size at 0 min and 120 min of induction (n > 300). (**D**) Diagram representing the coverage of StfZ and DicF antisense RNA on *ftsZ* sequence. The StfZ (blue line) covers the complete intergenic region and a significant portion of *ftsZ* 5′ region while DicF (brown line) partially covers the 5′ region.

### StfZ RNA influences FtsZ level and thereby cell division

Unlike studies in the case of *trans*-antisense RNAs (Majdalani et al., 2004; Papenfort et al., 2009; Guo et al., 2014), since the sequence of StfZ and its promoters are complementary to the reading frame of the target gene, *ftsZ*, which is essential for cell viability, generation of knockout mutant and/or promoter mutations could not be taken up to determine the physiological effect of the lack of expression of StfZ. Therefore, we tested the effects of StfZ overexpression on FtsZ protein level. A 423 bp region of *stfZ* was cloned and expressed from PAK12 (P_*lac-*_*stfZ*) strain. Strain PAK02 (vector control) and PAK12 (S1 Table) were induced with 1 mM IPTG for 2 hrs at 30°C and 42°C. A higher induction temperature of 42°C was used as per DicF *trans*-antisense RNA experiment, where the translation of FtsZ was blocked only at 42°C but not at lower temperatures (Tétart and Bouché, 1992). Upon induction, the levels of StfZ RNA increased to 180-fold and 600-fold, at 30°C and 42°C, respectively, compared to the level of the endogenous StfZ RNA in the pBS(KS) vector control cells (Fig. 4A). Consequentially, at 42°C, the level of FtsZ protein was 17% at 120 min post-induction of StfZ (Fig. 4B, upper panel, and C). The change in FtsZ level at 30°C was not significant. The coomassie blue stained PVDF membrane used for western blot analysis showed equal loading (Fig. 4B, lower panel). The *ftsZ* mRNA level, with the *ftsA* mRNA transcribed from the same operon as the control, did not change upon StfZ overinduction. Thus, the target sense RNA did not get degraded in the case of StfZ RNA-*ftsZ* mRNA interaction, unlike in many cases (Dühring et al., 2006; Giangrossi et al., 2010; Lee and Groisman, 2010; Bordoy and Chatterjee, 2015).

**Figure. 4.**
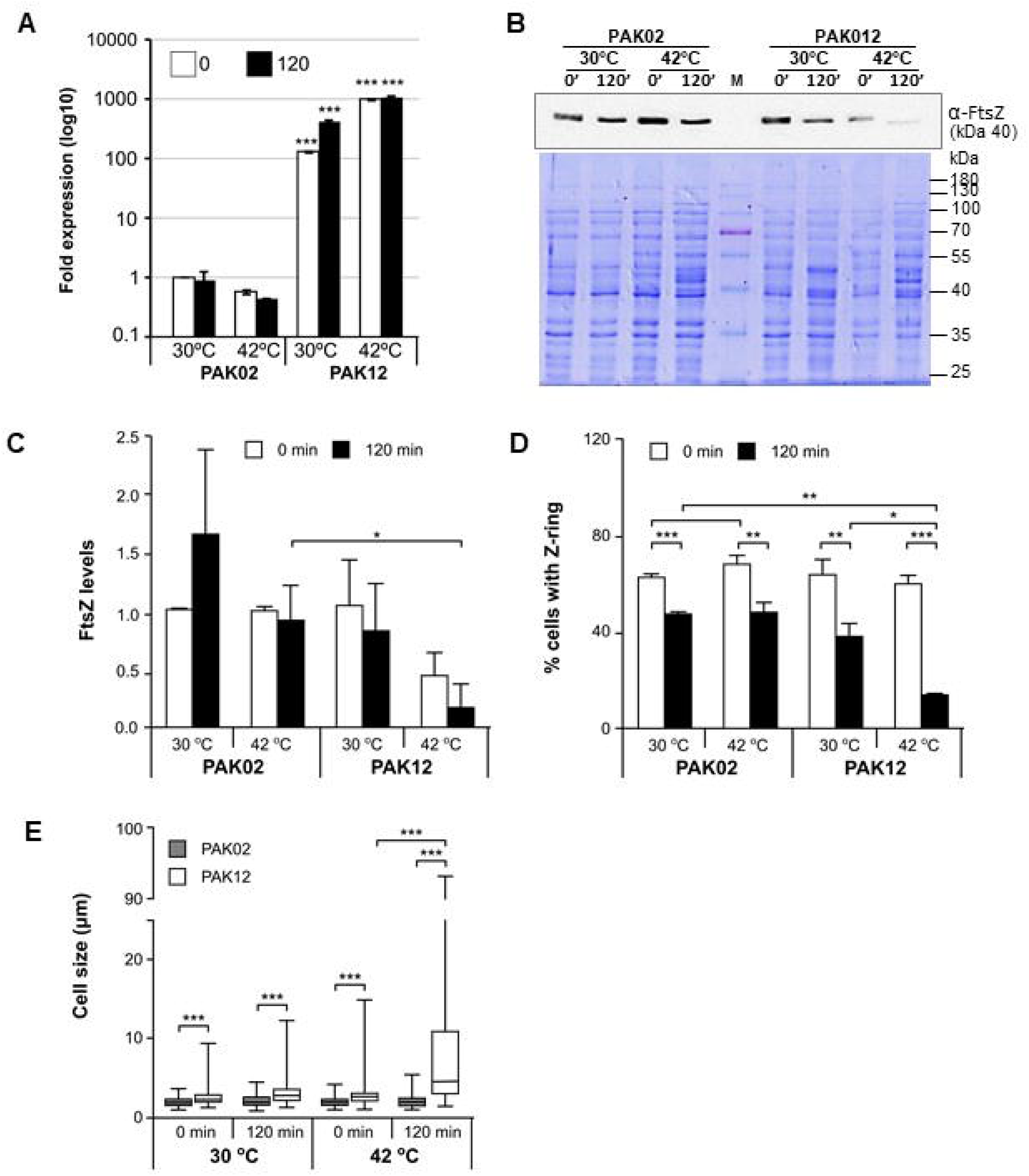
StfZ overinduction decreases FtsZ level and blocks cell-division. (**A**) Real time PCR for StfZ overexpression at 30°C and 42°C in vector control (PAK02) and StfZ expression strain (PAK12) (**B**) Western blot analysis for FtsZ level and equal loading profile of Coomassie-stained PVDF membrane used for the analysis. (**C**) Bar-graph for the quantitation of FtsZ using 200 μg total cell lysate after StfZ RNA induction for 0 min and 120 min at 30°C and 42°C. FtsZ level was calculated with respect to PAK02 sample from 0 min at 30°C. (**D**) Bar-graph for the reduction in the proportion of cells with FtsZ ring after StfZ overinduction. Cells were fixed after 0 min and 120 min of induction and immunostained for FtsZ. Cells with FtsZ ring were compared between 0 min and 120 min of induction. (**E**) Box-chart for cell-size measurement of PAK02 and PAK12 cells at 0 min and 120 min of induction at 30°C and 42°C. Statistical significance is indicated with asterisks (p < 0.05 *, p < 0.01 **, p < 0.001 ***). (**F**) Immunostaining for FtsZ in PAK02 and PAK12 cells from 30°C and 42°C. Left panel PAK02 (vector control) and right panel PAK12 (StfZ induction) at 0 min upper half panel and 120 min (lower half panel) are shown. Microscopy images are arranged in DIC, FtsZ-Cy3, DAPI stained nucleoid and merged from left to right in each panel.

The influence of reduced FtsZ level on cell division was examined by determining the proportion of cells with FtsZ rings by visualising immuno-stained FtsZ. At 0 min, ∼60-70% PAK12 population of contained FtsZ ring at 30°C and 42°C respectively (Fig. 4D). After 120 min of induction at 42°C, the population of cells with FtsZ ring reduced to ∼15% (Fig. 4D). Slightly elongated cells were noticed in the 120 min induced sample at 30°C, while the cells in the 120 min induced sample at 42°C were filamented (Fig. 4D, Fig 4E). The reduction in the proportion of cells with FtsZ ring and consequential cell elongation and/or filamentation correlated with the reduced FtsZ level (Fig. 4E). Induced expression of StfZ at 37°C for 120 min also showed about a 1.75-fold reduction in the FtsZ level as compared to the 0 min sample at 37°C (S1A Fig). Commensurate with this reduction, the proportion of cells with FtsZ ring also decreased significantly from ∼60% to ∼40% in 120 min (S1B Fig, S1C Fig).

Measurement of optical density of the cells carrying uninduced and induced StfZ RNA showed a significant increase in PAK02 and PAK12 mass (S1D Fig). However, PAK12 mass was significantly low at 120 min compared to that of PAK02, indicating inhibition of cell division and consequential lack of increase in cell number. The higher OD of PAK02 at 120 min might be from cell number increase due to cell division. Cfu at 120 min showed an increase in PAK02 population while the PAK12 cells did not show a significant increase at 120 min (S1E Fig). The cfu data corroborated the cell mass data that the lack of increase in cell mass in PAK12 was a result of lack of cell division while in PAK02, it was due to an increase in the cell number by cell division. SYTO9/PI staining confirmed that 120 min of StfZ induction did not affect the cell viability (S1F Fig). Taken together, the *cis*-encoded StfZ RNA emerges as a novel factor involved in the maintenance of FtsZ levels at different growth phases in *E. coli*.

## Discussion

### Role of StfZ RNA

The present study shows the complete sequence identity of StfZ RNA, with its 5′ and 3′ ends, promoters transcribing them and its ability to inhibit FtsZ translation. Higher StfZ levels imposes cell division block, resulting in cell elongation and filamentation. The effect on FtsZ protein reduction was not due to a decrease in the target *ftsZ* mRNA stability unlike in the case of many sense-antisense RNA interactions (Dühring et al., 2006; Giangrossi et al., 2010; Lee and Groisman, 2010; Bordoy and Chatterjee, 2015). The expression of StfZ throughout the entire growth phase at 1/6^th^ the proportion of *ftsZ* mRNA indicated stringent regulation of its expression commensurate with the levels of its target *ftsZ* mRNA. Such stringent regulation of the *cis*-antisense RNA, StfZ, to its sense *ftsZ* mRNA target might be to ensure that the concentration of *ftsZ* mRNA available for translation is always as per the growth phase-specific demand in the bacteria. The higher levels of both *ftsZ* mRNA and StfZ RNA synthesised during early phases of active growth and their decrease during late phases of growth are indicative of the ‘synthesis-as-per-demand’ mode of expression. We have not examined whether the level of StfZ RNA would change under any stress condition. Nevertheless, from the level of its expression being commensurate with the level of *ftsZ* mRNA at 1:6 ratio, it is tempting to speculate that under whatever stress condition FtsZ level would change, the level of StfZ RNA also would necessarily have to change accordingly to keep the available translatable *ftsZ* mRNA at the required level.

### Isoforms of StfZ RNA

Many *trans*-antisense RNAs have isoforms that are most often the processed forms of a primary transcript expressed from a single promoter. Some of such *trans*-antisense RNAs are the DicF (Bouché and Bouché, 1989), ArcZ (Papenfort et al., 2009; Soper et al., 2010), RprA (Majdalani et al., 2004), and MicL (Guo et al., 2014) RNAs of *E. coli*. In the case of StfZ RNA, the loss of activity of the -10 deletion mutants of the three promoters, predicted based on the 5′ end identification using PEA, validated the authenticity of the predicted promoters and the existence of three isoforms. The transcriptional activity of the three predicted promoters and its abrogation in Δ-10 construct indicates that the predicted promoter sequences are capable to initiating transcription therefore, PAE products are not a result of a processed product. However, bacterial promoters showing different activities at the various location, such as in the genomic context or as individual clones in a plasmid, have been reported in many instances (Cases and de Lorenzo, 2005; Davis et al., 2011; Hocine et al., 2015; Srivastava et al., 2016). Therefore, the presence of three potential PE products or three bands in the northern blot cannot alone conclude the existence of three promoters for StfZ RNA. *cis*-encoded nature of StfZ RNA to an essential gene does not permit their conclusive verification by mutating the promoters one at a time in the genome and checking for the decrease in the level of StfZ RNA and consequential increase in FtsZ level. Thus, studies of *cis*-encoded antisense RNAs have been possible only through overexpression, such as in the case of *ureB cis*-encoded antisense RNA against *ureAB* mRNA (Wen et al., 2011). Another example of a *cis*-encoded antisense RNA that exists in three isoforms is the *cis*-encoded GadY RNA, which regulates acid response genes in *E. coli* (Opdyke et al., 2004). All the three isoforms of GadY RNA are detected at all growth phases in a growth phase-dependent manner, like in the case of StfZ RNA (Opdyke et al., 2004).

It may be stated that differential expression of StfZ RNA from multiple promoters may help fine-tune the level of available *ftsZ* mRNA for translation, to avoid fluctuations in the FtsZ level (Ward and Lutkenhaus, 1985; Bi and Lutkenhaus, 1990). Such fine-tuning the of StfZ expression may be a logical necessity since the cells maintain a critical level of FtsZ, expressed from multiple promoters and through various other mechanisms (Vicente et al., 1998; Dewar and Dorazi, 2000). Further investigations centred on the regulation of expression of StfZ RNA and examination on the possibility of its interaction with antisense RNA binding proteins might reveal the mechanisms behind the antisense StfZ RNA mediated maintenance of the level of the essential cytokinetic protein, FtsZ, during cell division.

## Acknowledgements

The authors thank Dr. Raphael Valdivia for pFPV27 vector and Dr. William Margolin for pBAD33/*ftsZ*-*yfp*. Authors acknowledge technical help from Mrs. H. S. Rajeswari in immunostaining and western blotting and Dr. José Vicente Gomes for RNA-Seq data analysis.

## Author Contributions

Conceived and designed experiments: PA, DA, DS, RRN, KJ. performed experiments: DA, DS, RRN, KJ. Analysed data: PA, DA, DS, RRN, KJ, NM. Contributed reagents/materials/analysis tools: PA, DA, DS, RRN, KJ, NM. Wrote the manuscript: PA, DA, DS, RRN. The authors do not have any conflict of interest.

## List of Supporting Information

S1 Table: Bacterial strains used in the study. (DOC)

S2 Table: Plasmid vectors used in this study (DOC)

S3 Table: Reagents used in this study. (DOC)

S4 Table: Oligonucleotide used in the study. (DOC)

S1 Fig - FtsZ level, and cell division and viability status upon StfZ over induction. (PPT)

