## Supplementary figures and images for "Complete identity of the *cis*-asRNA StfZ and its influence on FtsZ protein level and cell division in *Escherichia coli*"

### S1 Fig

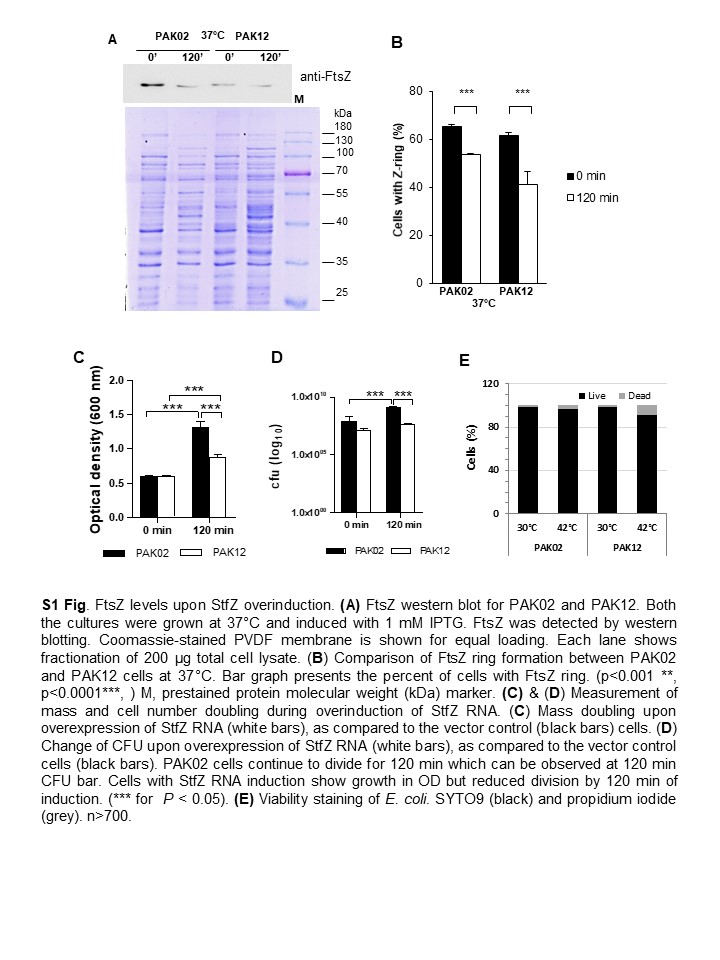
