## Supplementary material for "Complete identity of the *cis*-asRNA StfZ and its influence on FtsZ protein level and cell division in *Escherichia coli*": S1 Table

### S1 Table. List of bacterial strains used in this study

| STAIN | GENOTYPE | PLASMIDS | REFERENCE |
|---|---|---|---|
| *E. coli K12,* Wild type *(wt)* | ($F^+$ lambda+) | | (Blattner et al., 1997) |
| *E. coli JM109* | *recA1, supE44 endA1 hsdR17 gyr96 relA1 thiΔ(lac-proAB) F'[traD36proAB$^+$ lacI$^q$ lacZ ΔM15]* | | (Yanisch-Perron et al., 1985) |
| PAK-01 | K12, pBS(KS) | pBS(KS) | This study |
| PAK-02 | JM109, pBS(KS) | pBS(KS) | |
| PAK-03 | JM109, *stfZ* 3' RACE | pDA-01 | This study |
| PAK-04 | JM109, pFPV27 (mutgfp) | pFPV27 | This study |
| PAK-05 | JM109; *$P1_{stfZ}$-mutgfp* | pDA-02 | |
| PAK-06 | JM109; *P1*Δ-$10_{stfZ}$-*mutgfp* | pDA-03 | This study |
| PAK-07 | JM109; *$P2_{stfZ-}$mutgfp* | pDA-04 | This study |
| PAK-08 | JM109; *P2*Δ-$10_{stfZ}$-*mutgfp* | pDA-05 | This study |
| PAK-09 | JM109; *$P3_{stfZ}$*-P3-*mutgfp* | pDA-06 | This study |
| PAK-10 | JM109; *P3*Δ-$10_{stfZ}$-*mutgfp* | pDA-07 | This study |
| PAK-11 | JM109; *P1→$3_{stfZ}$-mutgfp* | pDA-08 | This study |
| PAK-12 | JM109; *$P_{lac}$-stfZ* | pDA-9 | This study |
| PAK-13 | JM109, pBAD-33-*ftsZ-yfp; $P_{lac}$-stfZ* | pDA-9, pBAD33-*ftsZyfp* | This study |
