## Supplementary material for "Complete identity of the *cis*-asRNA StfZ and its influence on FtsZ protein level and cell division in *Escherichia coli*": S2 Table

### S2 Table. List of plasmids used in this study

| PLASMID | DESCRIPTION | REFERENCE |
|---|---|---|
| pBS(KS) | Cloning & expression vector | Alting-Mees et al, 1989 |
| pFPV27 | Promoter probe vector, *mutgfp* for *E. coli,ColE ori, kan*$^R$ | Valdivia et al, 1996 |
| pDA1 | pBS(KS); *stfZ* 3' RACE, *amp*$^R$ | This study |
| pDA2 | Derivative of pFPV27, $P1_{stfZ}$*-mutgfp, kanR* | This study |
| pDA3 | Derivative of pFPV27, $P1\Delta\text{-}10_{stfZ}$*-mutgfp, kan*$^R$ | This study |
| pDA4 | Derivative of pFPV27, $P2_{stfZ}$*-mutgfp, kan*$^R$ | This study |
| pDA5 | Derivative of pFPV27, $P2\Delta\text{-}10_{stfZ}$*-mutgfp, kan*$^R$ | This study |
| pDA6 | Derivative of pFPV27, $P3_{stfZ}$-P3-*mutgfp, kan*$^R$ | This study |
| pDA7 | Derivative of pFPV27, $P3\Delta\text{-}10_{stfZ}$*-mutgfp, kan*$^R$ | This study |
| pDA8 | Derivative of pFPV27, $P1 \rightarrow 3_{stfZ}$*-mutgfp, kan*$^R$ | This study |
| pDA9 | Derivative of pBS(KS), $P_{lac}$*-stfZ, amp*$^R$ | This study |
| pBAD33-*ftsZyfp* | Derivative of pBAD33, $P_{\text{ara}}$-Ec-*ftsZyfp, amp*$^R$*,kan*$^R$ | W. Margolin |
