## Supplementary material for "Complete identity of the *cis*-asRNA StfZ and its influence on FtsZ protein level and cell division in *Escherichia coli*": S3 Table

### S3 Table. List of reagents used in this study

| Method | Buffer | composition |
|---|---|---|
| RNA Preparation | Lysis buffer | 100 mM Sodium acetate (pH 5.2), 10 mM EDTA (pH 8.0), 1% (w/v) SDS,100 mM NaCl, 100 mM β-mercaptoethanol, and 5 mM vanadyl ribonucleoside complexes |
| c-DNA preparation | Reaction buffer | 50 mM Tris-HCl (pH 8.3), 75 mM KCl, 3 mM $MgCl_2$, and 10 mM DTT |
| PNK reaction buffer | PNK 1x buffer | 50 mM Tris-HCl (pH 7.6), 10 mM $MgCl_2$, 5 mM DTT, and 0.1 mM spermidine |
| Northern hybridization | Pre-hybridization buffer | 7% SDS, 200 mM $Na_2HPO_4$ (pH 7.0) and 5 µg/ ml salmon sperm DNA |
| | Hybridization buffer | 7% SDS, 200 mM $Na_2$HPO4 (pH 7.0) and Biotin labeled probe |
| Immunostaining | PBS | 137 mM NaCl, 2.7 mM KCl, 10 mM $Na_2HPO_4$ (pH 7.4), 2 mM $KH_2PO_4$ |
| | PBST solution | 137 mM NaCl, 2.7 mM KCl, 10 mM $Na_2HPO_4$ (pH 7.4), 2 mM $KH_2PO_4$, Tween 20 (0.1%) |
| Western blot | PBST | 137 mM NaCl, 2.7 mM KCl, 10 mM $Na_2HPO_4$ (pH 7.4), 2 mM $KH_2PO_4$, Tween 20 (0.1%) |
| | blocking buffer | 5% w/v skimmed milk, 1x PBST |
