## Supplementary material for "Complete identity of the *cis*-asRNA StfZ and its influence on FtsZ protein level and cell division in *Escherichia coli*": S4 Table

**S4 Table. List of oligonucleotide used in the study**

| Oligo number | Oligo name | Sequence (5' to 3') | Comment |
|---|---|---|---|
| ODA-01 | EcZ-A/S-RT-f | GTTCCATTGGTTCAAACATAGTTTCTCTCC | RT-PCR |
| ODA-02 | EcZ-A/S-RT-r | CACATCTTAACGGTGAAGCTGAAGTAGAAA | RT-PCR, PEA |
| ODA-03 | Ec-16S-rRNA-RT-f | TGAAGACCGGCGGAAGAAG | RT-PCR |
| ODA-04 | Ec-16S-rRNA-RT-r | CACTTTATGAGGTCCGCTTGCT | RT-PCR |
| ODA-05 | EcftsZ-RT-f | TGAAGACCGGCGGAAGAAG | RT-PCR |
| ODA-06 | EcftsZ-RT-r | GAAGCAAATGCACGGATGGT | RT-PCR |
| ODA-07 | EcZ-A/S-PE2 | CCAATGGAACTTACCAATGACGCGG | PEA |
| ODA-08 | EcZ-A/S-RTr2 | CCTCAGGCGACAGGCACAAATCGGAGAGAAACT | PEA |
| ODA-09 | pFPV-27-mutgfp-RT-f | TCGGTTATGGTGTTCAATGCTT | RT-PCR |
| ODA-10 | pFPV27-mutgfp-RT-r | ACTTGACTTCAGCACGTGTCTTG | RT-PCR |
| ODA-11 | EcftsA-RT-f | CAAGGCGACGCACAGAAAAC | RT-PCR |
| ODA-12 | EcftsA-RT-r | TTCACCCCGCCTTTATCCA | RT-PCR |
| ODA-13 | Ec-ftsZ-A/S-locus-f | GGGGTACCTGTCTGCACCTTCCAGCGCC | PCR |
| ODA-14 | Ec-ftsZ-A/S-f | GGGTACCTTGGTGATACCGCTACCGATTTGAATCGTCTG | Cloning |
| ODA-15 | Ec-ftsZ-A/S-r | GCTCTAGATTTAACGGATTATGCTCAGGAGCCGTATTAT TCGA | Cloning |
| ODA-16 | EcZ-a/s-P1-f | GATCCCGCCGACGCCGATGACTTTAATCACCGCGTCATT GGTAAGTTCCATTGGTTCAAACATAGGGTAC | Cloning |
| ODA-17 | EcZ-a/s-P1-r | CCTATGTTTGAACCAATGGAACTTACCAATGACGCGGTG ATTAAAGTCATCGGCGTCGGCGG | Cloning |
| ODA-18 | EcZ-a/s-P2-f | GATCCGTCCAACCGCTGTTTTACGCAGCGCTTGTGCATC GGTATTTACCGCGAAGAATTCAACACGGTAC | Cloning |
| ODA-19 | EcZ-a/s-P2-r | CGTGTTGAATTCTTCGCGGTAAATACCGATGCACAAGCG CTGCGTAAAACAGCGGTTGGACG | Cloning |
| ODA-20 | EcZ-a/s-P3-f | GATCCCAGCCGCATTGCGGCCAACTTCTGGATTAGCGCC AGCGCCCAGTCCTTTGGTGATACCGCGGTAC | Cloning |
| ODA-21 | EcZ-a/s-P3-r | CGCGGTATCACCAAAGGACTGGGCGCTGGCGCTAATCCA GAAGTTGGCCGCAATGCGGCTGG | Cloning |
| ODA-22 | EcZ-a/s-P1-10f | GATCCCGCCGACGCCGATGACTTTAATCACCGCGTCATT GGCCATTGGTTCAAACATAGGGTAC | Cloning |
| ODA-23 | EcZ-a/s-P1-10r | CCTATGTTTGAACCAATGGCCAATGACGCGGTGATTAAA GTCATCGGCGTCGGCGG | Cloning |
| ODA-24 | EcZ-a/s-P2-10f | GATCCGTCCAACCGCTGTTTTACGCAGCGCTTGTGCATC GGCCGCGAAGAATTCAACACGGTAC | Cloning |
| ODA-25 | EcZ-a/s-P2-10r | CGTGTTGAATTCTTCGCGGCCGATGCACAAGCGCTGCGT AAAACAGCGGTTGGACG | Cloning |
| ODA-26 | EcZ-a/s-P3-10f | GATCCCAGCCGCATTGCGGCCAACTTCTGGATTAGCGCC AGCGCCTTTGGTGATACCGCGGTAC | Cloning |

| | | | |
|---|---|---|---|
| ODA-27 | EcZ-a/s-P3-10r | CGCGGTATCACCAAAGGCGCTGGCGCTAATCCAGAAGTT GGCCGCAATGCGGCTGG | Cloning |
| ODA-28 | EcZ-a/s-P3-T-f | CGGGATCCCAGCCGCATTGCGGC | Cloning |
| ODA-29 | EcZ-a/s-P1-T-r | GGGGTACCCTATGTTTGAACCAATGGAACTTACCAATG | Cloning |
| ODA-30 | 3'Lig-Prim-compl-KS-r | CGAGGTCGACGGTAGGTACCCC | 3’ RACE |
| ODA-31 | 3'Lig-Prim-KS-f | GGGGTACCTACCGTCGACCTCG | 3’ RACE |
